# Prolonged developmental programming of somatostatin neurons defines adolescent remodeling of the prelimbic, but not barrel, cortex

**DOI:** 10.64898/2026.07.27.740945

**Authors:** Avery R Sicher, Lauren N. Beloate, Varya Zlotnik, Keith Griffith, Erin Wolfanger, Hayreddin Said Ünsal, Nanyin Zhang, Nicole A. Crowley

## Abstract

Higher order association cortices like the prefrontal cortex (PFC) mature over a longer developmental window than sensory cortices, but whether genetically defined interneuron populations follow this region-wide delayed trajectory remains unclear. We compared the postnatal development of somatostatin-expressing interneurons in a higher-order cortical region (prelimbic cortex; PLC) and a sensory cortical region (somatosensory barrel cortex; S1BF) across adolescence using complementary structural, electrophysiological, and circuit analyses. We found that the maturation of SST neurons within these two cortical regions differs across time. S1BF SST neurons exhibited relatively linear maturation, whereas PLC SST neurons underwent continued dendritic remodeling, nonlinear intrinsic maturation, and age-dependent refinement of inhibitory output onto pyramidal neurons. Multivariate analyses likewise supported a more linear developmental trajectory in S1BF and a more prolonged, heterogeneous trajectory in PLC. This suggests divergent maturation of the PLC and S1BF and supports a broader, longer window of critical cellular plasticity in the PLC than in sensory cortices. Together, this reflects an extended developmental program at the level of a genetically defined inhibitory cell type. Because SST neurons are positioned to regulate pyramidal neuron output via control of dendritic integration, their delayed maturation is a crucial reflection of overall circuit function. These findings provide a cellular framework for the prolonged plasticity and developmental vulnerability known to be characteristic of adolescent prelimbic cortex.

**Significance statement:** Within the brain, higher order association cortex remains plastic long after sensory cortex has matured, but the cellular basis for this prolonged developmental window has remained unknown. We find that developmental timing is encoded within genetically defined microcircuits. Prelimbic somatostatin circuits continue to remodel after sensory somatostatin circuits have largely stabilized through changes in cell shape, action potential firing dynamics, and inhibitory signaling. This work points to a specific prelimbic inhibitory cell type that matures more slowly, and less linearly, than its counterparts in other brain regions, and may help explain why prelimbic circuits remain vulnerable during adolescence, representing a biological substrate for extended adolescent remodeling in prelimbic cortex, and potentially the overall extensive flexibility, but also cognitive susceptibility, seen during adolescence.

## Introduction

One of the defining features of mammalian association cortex (e.g., prefrontal cortical regions such as the prelimbic cortex) is its prolonged developmental trajectory. This delayed maturation is often thought to underlie why adolescence, the developmental period between childhood and adulthood characterized by physical, behavioral, and neurobiological changes, is a window of heightened vulnerability to stress, drug use, and the development of neuropsychiatric conditions (1–3). Developmental changes to the PFC in humans and rodents show improved efficiency of neurotransmission and refinement of overall activity through increases in neuromodulatory signaling, during which GABAergic systems undergo extensive remodeling (2–4). Although prolonged maturation of the PFC has been recognized for decades, it remains unclear whether this reflects a global delay across all neuronal populations or distinct developmental programs within specific circuit elements. In contrast to the PFC, sensory cortical regions including the somatosensory barrel cortex (S1BF) undergo comparatively rapid development within the first two postnatal weeks in rodents, coinciding with the structural formation of whisker barrels and the onset of active whisking (5, 6). This experience-dependent cortical maturation is well mapped in mammals (e.g., (7, 8)), with orderly development over a modest period of days. While the ‘rulebook’ of S1BF plasticity is incredibly-well characterized (9), a similar principled framework of PFC GABA neuron subtype development has not been undertaken.

Somatostatin (SST)-expressing neurons are the second largest subpopulation (20%) of GABAergic cells in the cortex (10), and are the earliest derived subpopulation of GABAergic cells. SST neurons begin to emerge in the medial ganglionic eminence during the second gestational week in mice (11), innervate superficial cortical layers shortly after birth (12, 13), and contribute to the formation of both excitatory and other inhibitory circuits in the first postnatal week (14–16). Few studies have probed how these neurons continue to mature. Although intrinsic properties of cortical SST neurons stabilize early in rodent development (13, 17, 18), other properties mediating action potential firing continue to mature into early adolescence in PFC SST neurons (19). In addition to controlling the activity of other GABAergic populations in the PFC (20–23), SST neurons are critical to pyramidal neuron inhibition via regulation of synaptic integration at distal dendrites (24–26), and this circuit within the PFC demonstrates unique, experience-dependent plasticity into adolescence (27). As PFC SST neurons are linked to neuropsychiatric conditions (28–31), a comprehensive understanding of the developmental trajectory of these neurons and their associated circuitry may therefore provide insight into the neurobiology of how these conditions emerge. If developmental programs differ across cortical region cellular subtypes, they may represent a previously unrecognized mechanism by which higher order association brain regions acquire their unique functional properties.

Limited studies assess cortical interneuron development in both juvenile and adolescent ages, and do not take comparative approaches across sensory and higher-order cortical regions, making it difficult to understand convergent and divergent forms of cortical development. If developmental programs differ across genetically defined subpopulations within cortical regions, they may contribute directly to regional specialization. In this study we explored this question by investigating morphological, electrophysiological, and circuitry changes in SST neurons in two cortical regions (PLC and S1BF) known to broadly undergo different developmental maturations. Our findings indicate that PLC SST neurons mature on a more prolonged timetable than S1BF SST neurons across morphology, intrinsic physiology, and GABAergic synaptic output activity. These results are consistent with the possibility that SST circuits contribute to extended adolescent remodeling in the PFC and reflect developmental programming at the level of a genetically defined inhibitory cell type.

## Results

### Protracted cellular growth and developmental pruning of SST neurons in the prelimbic, but not barrel, cortex occurs throughout adolescence

A hallmark and conserved developmental change in the adolescent PFC is the pruning of underutilized synaptic processes of pyramidal neurons (32–38), a process that has not been characterized in GABAergic populations. We characterized cellular characteristics and synaptic pruning of SST cells in the PFC and S1BF of SST-Cre:MORF3 mice in early (PND 28) and late (PND56) adolescence to quantify the extent of SST arborization at these ages (representative images and morphological tracing from both regions, **Figure 1A-B**). In S1BF, a 2- way mixed-effects ANOVA (factors: age, radius from soma) revealed a significant effect of radius from the soma on the number of dendritic intersections (*F*_radius from soma_(3.702, 92.55) = 40.31, *p <* 0.0001), but no effect of age (*F*_age_(1, 25) = 0.9256, *p* = 0.3452) nor an interaction (*F*_radius x age_(3.702, 92.55) = 0.8019, *p* = 0.5187; **Figure 1C**). In contrast to the lack of morphological changes seen in S1BF SST neurons, we observed a significant difference in the number of dendritic intersections as a function of age in PLC SST neurons (**Figure 1D**; *F*_radius x age_(3.955, 75.14) = 4.004, *p* = 0.0055). In S1BF, there was not a significant difference in the area under the curve of the radius x dendritic intersections graph (**Figure 1E**; *t*(24.09) = 1.592, *p* = 0.1243) indicating no change in the cumulative intersections of S1BF SST neuronal processes. Similarly, we did not detect age-specific changes in soma morphology of S1BF SST neurons across parameters including soma area (*t*(24.82) = 1.708, *p* = 0.1000), perimeter (*t*(24.98) = 1.657, *p* = 0.1101), and circularity (*t*(22.10) = 0.1298, *p* = 0.8979), as indicated by unpaired t-tests with Welch’s correction (**Figure 1F,I,J**).

**Figure 1.**
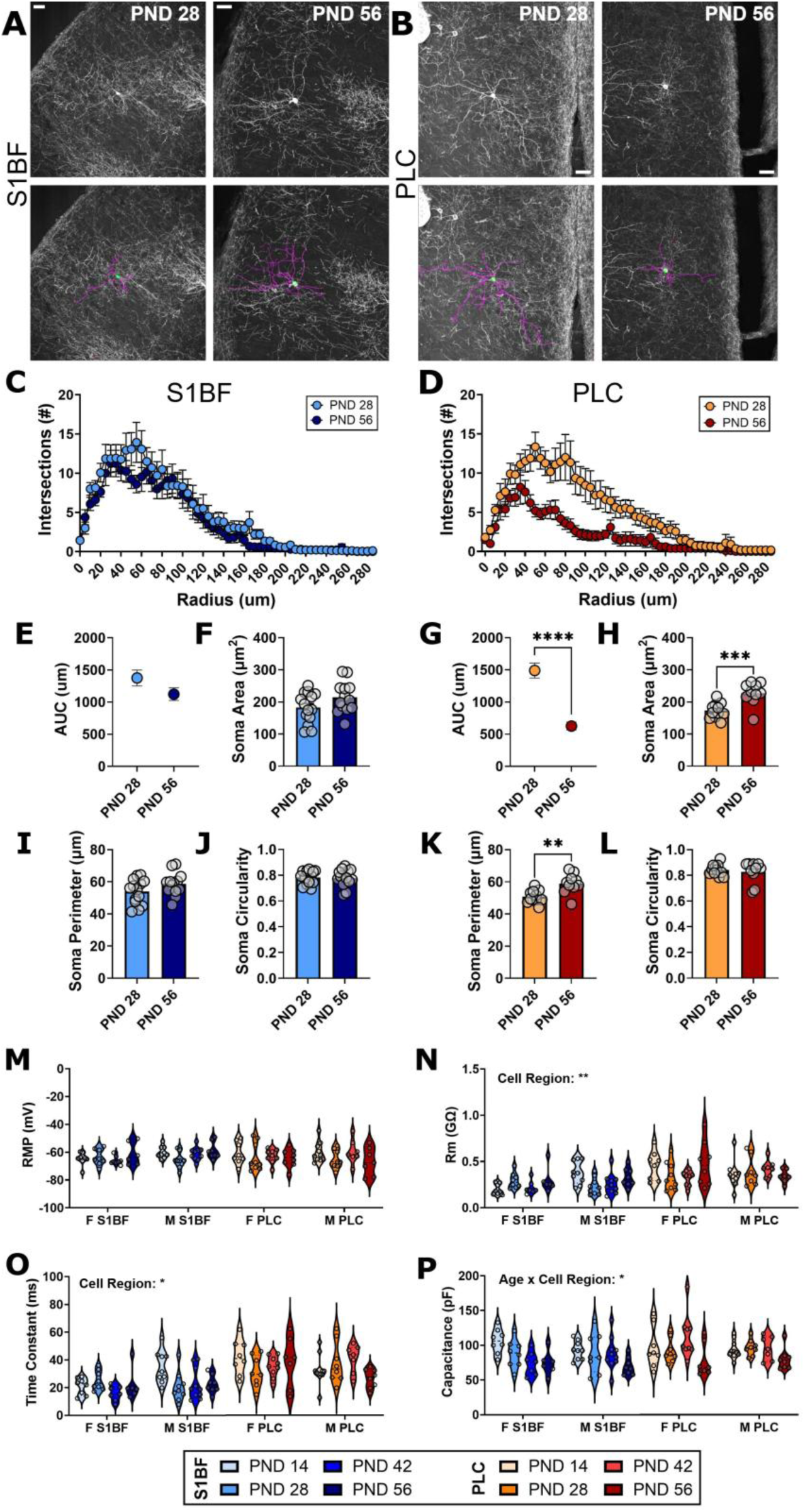
PFC SST neurons undergo protracted dendritic pruning compared to S1B1 SST neurons, despite few age-related changes in passive membrane properties. A-B) Representative images (top) and tracings (bottom) of SST neurons in S1BF and PLC at PND 28 and PND56. (C-D) Dendritic complexity is reduced in PL SST neurons during adolescence with no changes in S1BF SST neurons. (E-L) Soma morphology properties show protracted development in PL but not S1BF SST neurons. Morphology *n:* S1BF: PND28 = 14 cells, 9 mice; PND56 = 13 cells, 8 mice. PLC *n:* PND28 = 11 cells, 7 mice; PND56 = 10 cells, 6 mice. (M-P) Passive membrane properties of S1BF and PL SST neurons show region specificity.

In contrast to the S1BF, the area under the curve of the radius x dendritic intersections for PLC SST neurons was significantly higher at PND 28 than at PND 56 (*t*(15.09) = 6.498, *p* < 0.0001), indicating an overall reduction in the number of intersections across radii in older mice. An unpaired t-test with Welch’s correction showed that the soma area (**Figure 1H**; *t*(adjusted df=15.69) = 4.140, *p* = 0.0008) and soma perimeter (**Figure 1K**; *t*(15.05) = 3.567, *p* = 0.0028) of PLC SST neurons increased from PND 28 to PND 56, while the soma circularity remained unchanged (**Figure 1L**; *t*(13.32) = 0.5308, *p* = 0.6043).

We next explored electrophysiological markers of cell maturity in SST-Cre:Ai9 mice. Basal electrophysiological properties of SST cells showed a variety of region-specific and age by region interactions. GLME (data presented as Coefficient Estimate ± SEM) did not indicate that age (−0.076802 ± 0.081836, *p =* 0.3498), cell region (−6.0416 ± 4.3674, *p* = 0.16903), sex (−1.2778 ± 4.3613, *p* =0.77002), nor any interactions were significant predictors of RMP (**Figure 1M**). Cell region was a significant predictor of membrane resistance (**Figure 1N**; -0.21894 ± 0.083597; *p* = 0.009911), but not age (0.00015918 ± 0.001551; *p =* 0.91842), or sex (− 0.025279 ± 0.082795, *p =* 0.76063). Cell region was a significant predictor of membrane time constant (**Figure 1O**; -16.406 ± 7.0149; *p* = 0.020945) but not age (−0.062518 ± 0.12836; *p =* 0.62707), sex (−2.3906 ± 6.8785; *p =* 0.72877), nor any interactions. For membrane capacitance, the interaction between age and cell region was a significant predictor (**Figure 1P**; -0.80828 ± 0.36399; *p =* 0.028181). These results show that significant morphological changes occur in SST neurons in the PLC only, while passive membrane properties show variation across these two cortical regions.

### Intrinsic electrophysiological properties of S1BF SST neurons undergo rapid development, while PLC SST neurons have a protracted maturation

Despite evidence that passive membrane properties of frontal SST neurons stabilize after the first two postnatal weeks, other properties of intrinsic excitability continue to mature in frontal SST neurons past this timepoint (19). In contrast to frontal cortex, somatosensory SST neurons develop rapidly after the onset of active whisking (17, 39). However, studies have not directly compared how SST excitability develops across cortical regions.

We observed several developmental changes in SST excitability, many of which were region-dependent (representative traces in **Figure 2A-B**). Cell region was a significant predictor of rheobase (**Figure 2C**; 42.859 ± 13.983; *p =* 0.0026672), but not age (0.2912 ± 0.24739; *p =* 0.24141), sex (14.13 ± 13.297; *p =* 0.2900), nor any interactions. Action potential threshold was not significantly predicted by cell region (**Figure 2D**; -0.74798 ± 3.3026; *p =* 0.82119), age (0.054408 ± 0.058807; *p =* 0.35665), or sex (5.7912 ± 3.1587; *p =* 0.069123), nor any interactions. The interaction between age and sex (**Figure 2E**; -0.011577 ± 0.0046868; *p =* 0.014859) as well as cell region (−0.34614 ± 0.15785; *p =* 0.030172) were significant predictors of action potential half-width, indicating differences in maturation in the speed at which neurons can fire. Afterdepolarization (ADP) was significantly predicted by the interaction between cell region and sex (**Figure 2F**; -3.2269 ± 1.5325; *p =* 0.037234) and by age (0.072842 ± 0.020855; *p =* 0.00066173). Fast afterhyperpolarization (fAHP) did not have any significant predictors, including cell region (**Figure 2G**; -1.9469 ± 2.6501; *p =* 0.46392), age (−0.031479 ± 0.050707; *p =* 0.53586), sex (−3.8211 ± 2.6928; *p =* 0.15838), or interactions between these variables. Both cell region (**Figure 2H**; 0.025868 ± 0.0095837; *p =* 0.0079136) and age (0.00052679 ± 0.00016955; *p =* 0.0023403) were significant predictors of sag ratio.

**Figure 2.**
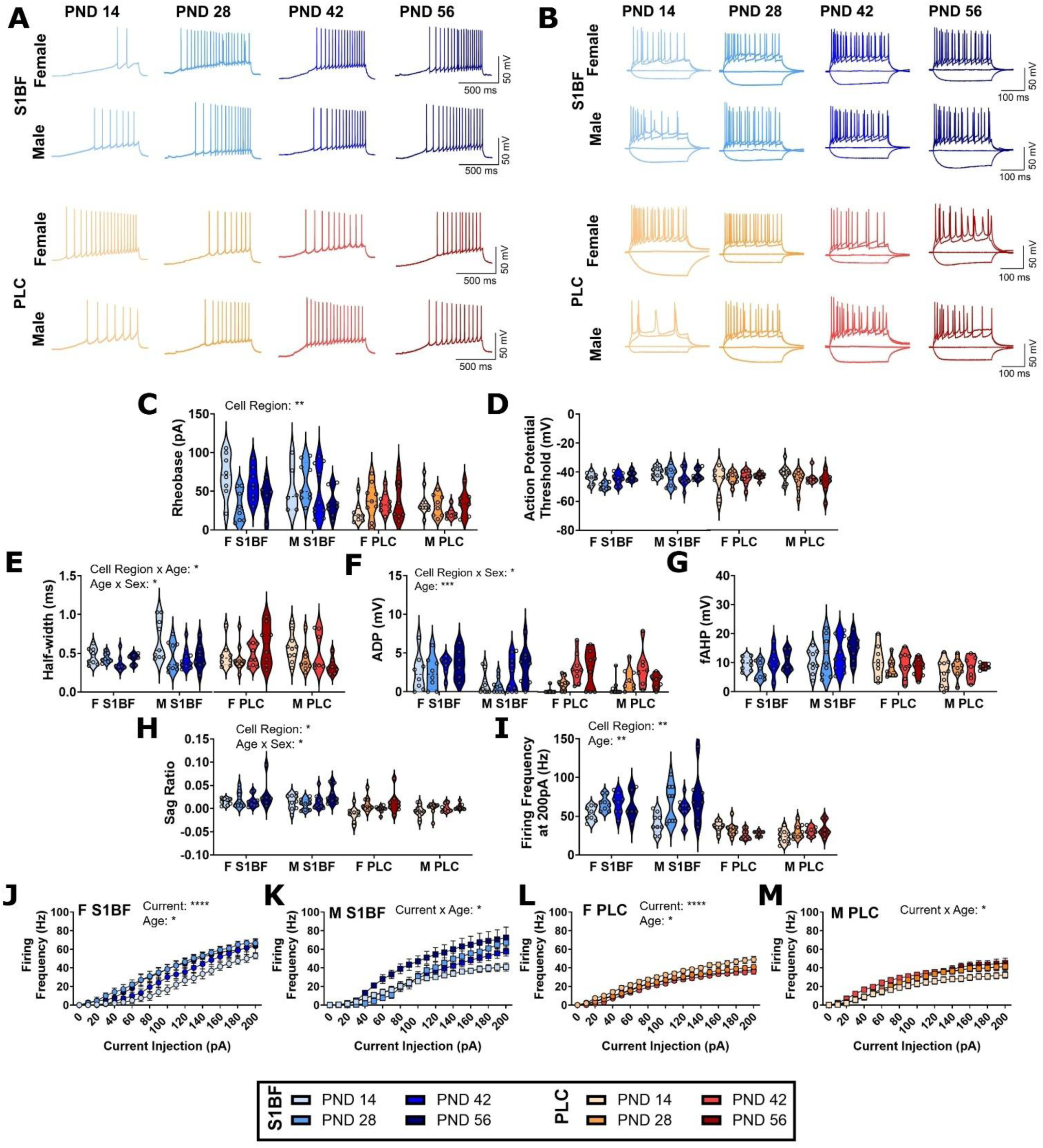
Measures of intrinsic excitability of cortical SST neurons across early postnatal development. (A-B) Representative traces of rheobase and current-induced firing experiments. (C) Rheobase is significantly different across cell regions. (D) No changes in action potential threshold due to age, cell region, or sex.(E) Action potential half-width development across cell region and sex. (F) Afterdepolarization emerges across postnatal age and is altered by the interaction between cell region and sex. (G) No changes in fast afterhyperpolarization due to age, cell region, or sex. (H) Sag differs across cell region and due to an age x sex interaction. (I) Current-induced firing after a 200pA current injection significantly varies based on cell region and age. (J-L) Current-induced firing across all depolarizing current injections split by cell region and sex.

Finally, peak firing frequency, taken from the 200pA current step, was significantly predicted by interactions between cell region and age (**Figure 2I**; 0.57766 ± 0.25707; *p =* 0.026388) and between age and sex (0.56007 ± 0.24881; *p =* 0.02613). We followed up on these interactions by assessing current-induced action potential firing within SST neurons of each cell region and sex using a 2-way ANOVA for each group (factors: age, current injection step). For SST neurons in S1BF of female mice, there were significant main effects of current (**Figure 2J**; *F_current_* (2.324, 60.43) = 308.5; *p <* 0.0001) and age (*F_age_* (3, 26) = 3.866; *p* = 0.0207). In S1BF of male mice, there was a significant interaction between current and age (**Figure 2K**; *F_current x age_* (5.106, 54.47) = 3.129; *p* = 0.0143). In S1BF of both sexes, current-induced firing increased after PND14. For SST neurons in the PLC of female mice, there was an expected main effect of current (**Figure 2L**; *F_current_* (2.148, 62.30) = 305.1; *p* < 0.0001) and a main effect of age (*F_age_* (3, 29) = 4.131; *p* = 0.0148). In the PLC of male mice, there was a significant interaction between current and age (*F_current x age_* (5.972, 55.73) = 2.463; *p* = 0.0351).

We repeated our intrinsic excitability protocols while holding cells at a common membrane potential of -70mV (**Supplemental Figure 1**). The interaction between cell region, age, and sex was a significant predictor of rheobase at -70mV (**Supplemental Figure 1A**; 2.0469 ± 0.78141; *p =* 0.0099142). The interaction between age and sex was a significant predictor of action potential threshold at -70mV (**Supplemental Figure 1B**; - 0.21255 ± 0.093631; *p =* 0.024944). The interaction between cell region and age was a significant predictor of the firing frequency during the 200pA current step at -70mV (**Supplemental Figure 1C**; 0.7447 ± 0.27899; *p =* 0.0086191). We followed up on this interaction using 2-way ANOVAs to assess current-induced firing within each sex and cell region group. For SST neurons in S1BF of female mice, there was a significant current x age interaction (**Supplemental Figure 1D**; *F_current x age_* (5.000, 45.00) = 3.120; *p* = 0.0168). For SST neurons in S1BF of male mice, there was a significant effect of current (**Supplemental Figure 1E**; *F_current_* (2.302, 73.66) = 137.0; *p <* 0.0001) but not age (*F_age_* (3, 32) = 0.9174; *p =* 0.4435), nor an interaction (*F_current x age_* (6.906, 73.66) = 1.883; *p* = 0.0854). In the PLC of female mice, there was a significant effect of current (**Supplemental Figure 1F**; *F_current_* (2.488, 72.16) = 284.4; *p* < 0.0001) but not an effect of age (*F_age_* (3, 29) = 1.959; *p* = 0.1423) nor an interaction (*F_current x age_* (7.465, 72.16) = 0.9169; *p =* 0.5030). Finally, in the PLC of male mice, there was a significant effect of current (**Supplemental Figure 1G**; *F_current_* (1.603, 44.89) = 209.9; *p* < 0.0001) but no effect of age (*F_age_* (3, 28) = 2.346; *p* = 0.0942) or interaction (*F_current x age_* (4.810, 44.89) = 1.539; *p* = 0.1989). Together, these findings show that the intrinsic excitability as well as action potential properties of SST neurons are developmentally regulated, but not uniformly across cortical regions.

### Electrophysiological properties of S1BF SST neurons develop linearly with age, while PFC shows less predictability

To further understand how intrinsic properties of SST neurons vary as a function of age, sex, and cortical region, we used correlations and Principal Component Analysis (PCA; **Figure 3**). In S1BF, age was significantly linearly correlated with peak firing frequency (**Figure 3A**; *r*(66) = 0.336, *p* = 0.006), membrane time constant (*r*(66) = -0.246, *p* = 0.047), membrane capacitance (*r*(66) = -0.458, *p* = 0.000123), action potential half-width (*r*(66) = -0.303, *p* = 0.014), afterdepolarization (*r*(66) = 0.354, *p* = 0.004), and voltage sag (*r*(66) = 0.274, *p* = 0.026). In contrast, in the PLC, only afterdepolarization (**Figure 3B**; *r*(66) = 0.449, *p =* 0.0001565) and voltage sag (*r*(66) = 0.349, *p* = 0.004) were significantly correlated with postnatal age. These findings show a linear relationship between age and intrinsic properties in SST neurons of S1BF, suggesting a more stereotyped development than SST neurons of the PLC. Radar plots were created to further visualize changes in passive (**Figure 3C**) and active (**Figure 3D**) properties of SST neurons across development. For these plots, data from each sex x region group were normalized by z-score to the average of all collected cells. Z-scores for rheobase, action potential threshold, and membrane time constant were inverted so that more positive values were reflective of increased excitability.

**Figure 3.**
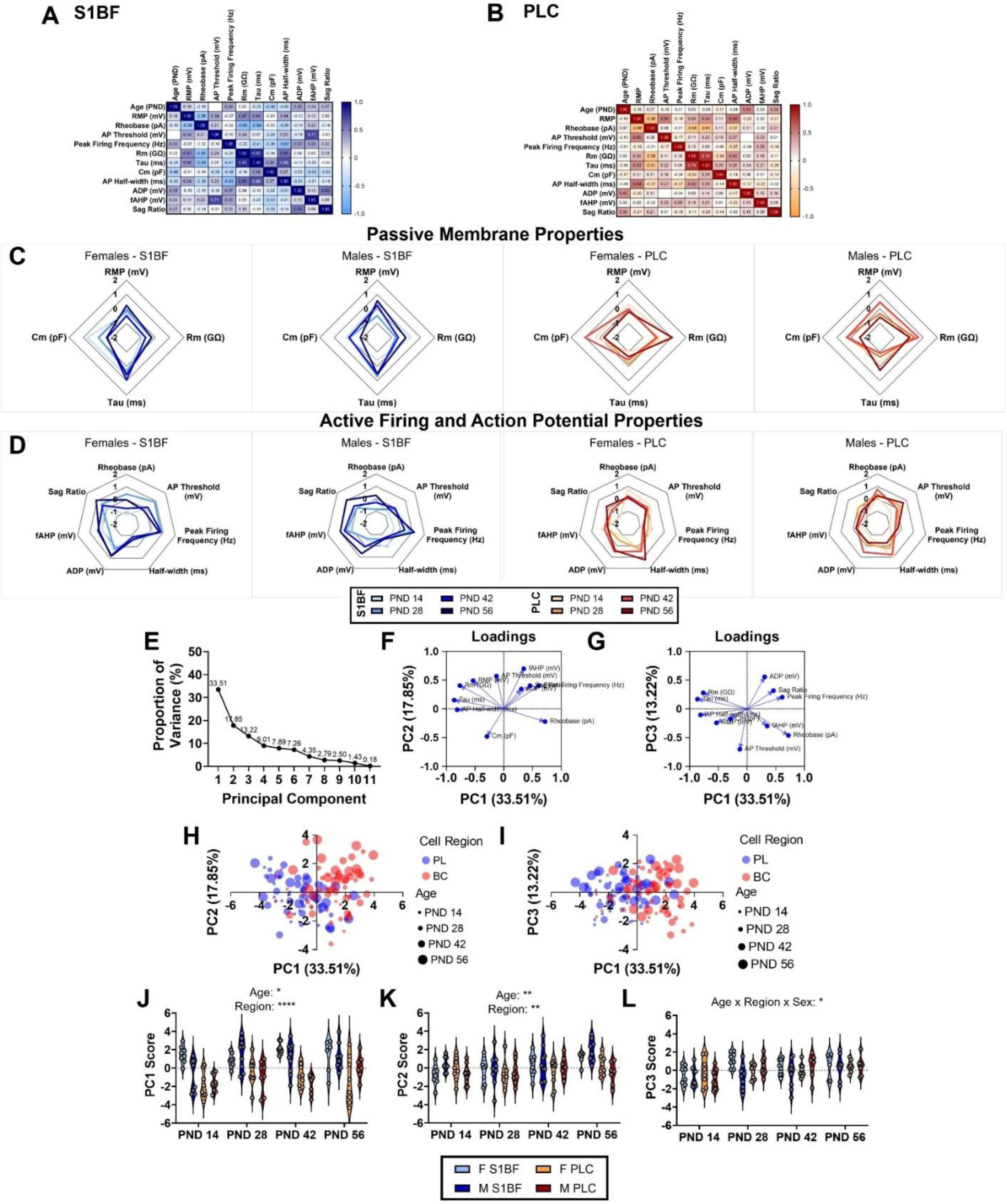
Membrane property analysis shows linear postnatal development of S1BF, but not PLC, SST neurons. (A-B) Pearson correlation matrices of extracted membrane properties show that postnatal age correlates with many properties in S1BF neurons. In PLC neurons, age correlates with the emergence of afterdepolarization and voltage sag. (C-D) Radar plots of passive and active membrane properties of SST neurons separated by sex and region. (E) Proportion of variance explained by 11 principal components (PCs) identified by principal component analysis. PCs 1-3, explaining about 65% of the variance, were selected by parallel analysis. (F-G) Variable loadings for PCs 1 and 2 (F) and PCs 1 and 3 (G). (H-I) Scores, representing individual cells, for PCs 1 and 2 (H) and PCs 1 and 3 (I). The color of the score value corresponds to the region and the size represents the age of the mouse from which the cell was recorded. (J-L) PC scores as a function of age for each sex x region group.

Given the number of electrophysiological properties measured, principal component analysis (PCA) was used to identify patterns in the intrinsic properties of SST neurons across postnatal development. PCA with parallel analysis identified 3 principal components accounting for about 65% of the variation in the data (**Figure 3E-L**). PC1 accounted for 33.51% of the variation in the data (**Figure 3F**) and had the greatest loading scores from membrane time constant (−0.864), action potential half-width (−0.811), and membrane resistance (−0.764). PC2 accounted for 17.85% of the variation and had the greatest loading scores from fast afterhyperpolarization (0.698), action potential threshold (0.569), and resting membrane potential (0.490). Finally, PC3 accounted for 13.22% of the variance in the data (**Figure 3G**) and had the largest loading values from action potential threshold (−0.703), afterdepolarization (0.556), and rheobase (−0.462). Scores from each cell are represented, visualized by sex and by cell region, in **Figure 3H-I**. We used 3-way ANOVA (factors: age, sex, cell region) to see if PC scores were altered across development, by cell region, or by sex of the mouse. There was a significant main effect of region (**Figure 3J**; *F_cell region_* (1, 115) = 70.42; *p <* 0.0001) and of age (*F_age_* (3, 115) = 3.380; *p =* 0.0207), but no main effect of sex (*F_sex_* (1, 115) = 1.174; *p =* 0.2809) nor any interactions on PC1 score. Similarly, there were significant main effects of age (**Figure 3K**; *F_age_* (3, 115) = 4.175; *p =* 0.0076) and of region (*F_cell region_* (1, 115) = 9.690; *p =* 0.0023) on PC2 score, but not sex (*F_sex_* (1, 115) = 0.6171; *p =* 0.4337), nor any interactions. Finally, for PC3 scores, there was a significant interaction between age, cell region, and sex (**Figure 3L**; *F_age_*(3, 115) = 2.974; *p =* 0.0346). The findings here, across multiple integrative analyses, show that intrinsic properties of SST neurons vary across postnatal development and across cortical regions.

### Developmental changes in SST-mediated signaling onto pyramidal neurons in PL, but not S1BF cortex

To determine whether changes in synaptic communication between SST neurons and their primary target (pyramidal neurons) occur during postnatal development, we performed optogenetic circuit mapping (representative traces in **Figure 4A**). SST neurons in both PLC and S1BF fired reliably in response to 470nm stimulation (**Supplementary Figure 2**). In S1BF pyramidal neurons recorded from female mice, we found an expected significant main effect of interstimulus interval (ISI) on paired pulse ratio (individual datapoints from female mice across ISIs shown in **Supplementa**l **Figure 3A**; **Figure 4B**; *F_ISI_* (2.002, 64.06) = 42.68; *p* < 0.0001) but no effect of age (*F_age_* (3, 32) = 0.7279; *p* = 0.5429) nor an interaction (*F_age x ISI_* (6.005, 64.06) = 1.732; *p* = 0.1278). We observed similar results in S1BF pyramidal neurons from male mice, where there was a significant main effect of ISI (individual datapoints from male mice across ISIs shown in **Supplementa**l **Figure 3B**; **Figure 4C**; *F_ISI_* (2.671, 85.49) = 72.67; *p* < 0.0001) but not age (*F_age_* (3, 32) = 0.7882; *p =* 0.5094) nor an interaction (*F_age x ISI_* (8.014, 85.49) = 0.8674; *p* = 0.5473).

**Figure 4.**
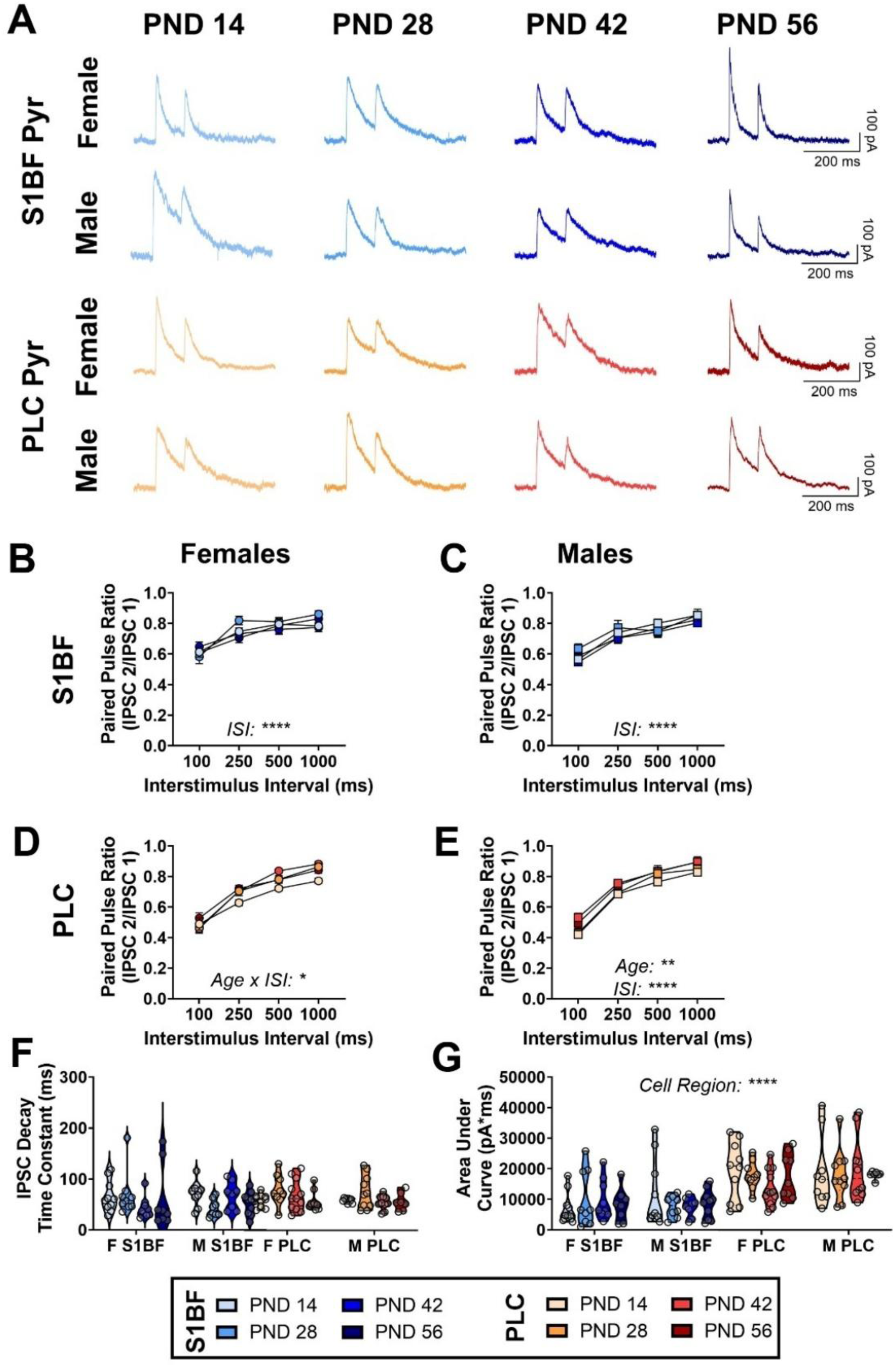
Optogenetic circuit mapping shows age-related remodeling at SST->Pyr synapses in the PLC, but not S1BF. (A) Representative traces of paired pulse recordings onto pyramidal neurons in S1BF and PLC. (B) No effect of age on paired pulse ratio (PPR) in S1BF pyramidal neurons of female mice. (C) Age did not affect PPR in S1BF pyramidal neurons from male mice. (D) There was a significant interaction between age and interstimulus interval on PPR in PLC pyramidal neurons of female mice. (E) Significant main effects of age and interstimulus interval on PPR of PLC pyramidal neurons from male mice. (F) No effect of age, cell region, or sex on decay time constant of the first evoked IPSC. (G) Area of the evoked IPSC varied across cell regions. IPSC: inhibitory postsynaptic current.

In PL pyramidal neurons from female mice, we found a significant interaction between age and interstimulus interval (individual datapoints from female mice across ISIs shown in **Supplemental Figure 4A**; **Figure 4D**; *F_age x ISI_* (5.143, 61.71) = 2.362; *p* = 0.0486). In PL pyramidal neurons recorded from male mice, we found a significant main effect of age (individual datapoints from male mice across ISIs shown in **Supplemental Figure 4B**; **Figure 4E**; *F_age_* (3, 32) = 5.943; *p* = 0.0024) in addition to the expected main effect of ISI (*F_ISI_* (1.624, 51.96) = 275.7; *p* < 0.0001). There was no significant main effect of cell region (**Figure 4F**; *F_region_*(1, 132) = 0.8781; *p* = 0.3504), age (*F_age_* (3, 132) = 1.006; *p* = 0.3922), sex (*F_sex_*(1, 132) = 0.3534; *p* = 0.5532), nor any interactions, on the IPSC decay time constant, measured from the first IPSC at the 1000ms ISI. A 3-way ANOVA (factors: cell region, age, sex) indicated that the area under the first IPSC of the 1000ms ISI varied significantly by region (**Figure 4G**; *F_cell region_* (1, 132) = 55.51; *p* < 0.0001) but not age (*F_age_*(3, 132) = 0.3140; *p* = 0.8152) or sex (*F_sex_*(1, 132) = 0.4754; *p* = 0.4917), nor any interactions. Together, these findings suggest that SST-Pyr synapses undergo developmental regulation in the PL cortex but not S1BF.

## Discussion

Our findings demonstrate that developmental timing of cortical functioning is encoded within genetically defined inhibitory microcircuits. Whereas sensory cortical (S1BF) SST neurons largely complete maturation before adolescence, homologous neurons in prefrontal cortex (PFC) continue structural, intrinsic, and synaptic remodeling throughout this period, identifying prolonged maturation of prefrontal SST interneurons as a cellular substrate for extended association-cortex plasticity. This further suggests that the timing of cellular development itself is a specialized feature of association cortex, providing a cellular basis for both the adaptability of adolescent cognition and the heightened vulnerability of the PFC during this critical developmental period.

### Continued morphological remodeling identifies an extended developmental program in PLC SST neurons

While S1BF SST neurons had largely stabilized early in development in our experiments, PLC SST neurons continue to undergo substantial structural remodeling throughout adolescence, characterized by increasing soma size and dendritic pruning. Although several studies have reported similar patterns of pruning in PFC pyramidal neurons, to our knowledge, this is the first study to characterize morphology changes in a subtype of GABAergic neurons during adolescence. We did not observe age-related morphology changes in S1BF SST neurons, consistent with results from others in other sensory cortex (40). This provides the first comprehensive evidence that PFC SST neurons retain plasticity further into development than previously thought. The protracted dendritic remodeling seen in PLC SST neurons is consistent with the continuing maturation of function of these cells, and suggests the need for a longer developmental period for glutamatergic inputs onto SST neurons to mature and for their dendritic arbors to stabilize than observed in other cortical regions (13). Because SST neurons integrate excitatory inputs via their dendritic arbor, continued dendritic remodeling is expected to alter how these neurons respond to these still maturing excitatory inputs. Complementary changes in neuronal soma size can reflect increased cellular complexity, and potentially development of the population’s ability to encode information (41). Together, these changes point towards ongoing maturity of the SST population within the PLC, in contrast to an earlier stabilized S1BF population.

### Intrinsic maturation reflects functional refinement of PLC SST neurons

Overall, we find that while passive membrane properties stabilized relatively early in both cortical regions, intrinsic excitability continued to mature in fundamentally different ways. In particular, PLC SST neurons became progressively less excitable despite continued maturation of intrinsic electrophysiological properties, suggesting that developmental refinement involves specialization of neuronal function rather than simply increasing excitability.

Our findings are consistent with previous reports that passive membrane properties of PFC (13, 18) and S1BF (17) SST neurons are largely stable after the second postnatal week. Interestingly, we saw a variety of differences in properties of SST neurons across cortical regions – neurons from S1BF tended to be more excitable than those in the PLC, as measured by current-induced firing and peak firing frequency. We also observed the relatively late emergence of some action potential properties, including afterdepolarization, in SST neurons from both regions. The emergence of afterdepolarization indicates continued maturation of intrinsic membrane properties beyond the developmental period during which passive properties have stabilized. Because afterdepolarization is characteristic of mature Martinotti neurons (42, 43), these findings suggest that maturation of SST neurons extends well beyond early postnatal development. However, we saw an age-related decrease in current-induced firing in PLC SST neurons from female mice, despite the emergence of afterdepolarization, suggesting that appropriate developmental maturation of this circuit does not simply require increased cellular intrinsic excitability, but perhaps also more selective cell-cell communication.

We performed several additional analyses to integrate changes across intrinsic measurements to identify broader patterns in SST development across cortical regions. Across multiple electrophysiological measurements, our PCA suggests that rather than individual electrophysiological properties maturing independently, there are distinct coordinated developmental programs across cortical regions. First, we observed that age significantly correlated with multiple properties in S1BF, but not PLC, SST neurons. We interpret this as evidence that S1BF SST neurons undergo a stepwise, progressed maturation in contrast to PLC SST neurons, which only showed linear emergence of a small number of properties (afterdepolarization and voltage sag). The orderly progression observed in S1BF contrasts sharply with the heterogeneous trajectories observed in PLC, supporting the idea that higher-order association cortex maintains multiple developmental states across adolescence.

There are a few potential interpretations of our overall findings regarding SST excitability in the PFC. One is that mature inhibitory neurons are capable of processing greater input specificity, where neuronal firing relies more on network-driven activation (rather than intrinsic mechanisms of activation) as cortical circuits become more refined. This interpretation suggests that reduced intrinsic excitability would reflect an accompanying stronger synaptic recruitment, which was not assessed in the current study, but is supported by the overall refinement of dendritic arborization. An alternative potential interpretation is that proper development involves a shift from broadly responsive SST neurons to more selectively recruited SST neurons during adulthood. Future would can explore the real-time recruitment of genetically-defined populations of SST neurons across development during *in vivo* tasks to understand shifts in the pool of coordinated SST neurons.

### Continued refinement of PLC SST GABAergic activity further suggests prolonged maturation of prefrontal inhibitory circuits

Our optogenetic circuit mapping experiments extend our cell morphology and excitability findings beyond intrinsic maturation by demonstrating that developmental remodeling also persists at the level of inhibitory synaptic output. Whereas SST-mediated inhibition of pyramidal neurons remained stable across development in S1BF, synaptic properties continued to evolve in PLC, indicating that maturation of PFC SST neurons extends to influence its downstream targets (and thus overall PLC output activity). In our optogenetic circuit mapping experiments, we saw that age modulated PPR of SST-mediated IPSCs onto pyramidal neurons only in the PLC. This is consistent with a previous study in the infralimbic PFC which found that SST inhibition of pyramidal neurons was highest in adolescence and sensitive to experience with a fear conditioning assay (27). Because there were no other age-dependent effects on other properties of the IPSC (potentially due to differences in GABA_A_ receptor subunit expression) we interpret these changes as differences in the readily releasable pool of GABA in PLC SST neurons across development, compared to a more stable pool in SIBF. An alternative, though not mutually exclusive, explanation is that developmental changes in SST neuron number may alter the population of channelrhodopsin-expressing neurons contributing to the recorded responses, as SST cell density in PFC follows an inverted U-shaped developmental trajectory (44).

Because SST neurons preferentially target pyramidal neuron dendrites, continued refinement of SST output is expected to influence dendritic integration, cortical gain control, and the balance between excitation and inhibition within prefrontal microcircuits. Continued maturation of these synapses may therefore contribute to the gradual stabilization of cortical computation that accompanies the transition from adolescence to adulthood.

### Extended maturation of SST neurons may shape adolescent susceptibility to experience

Our findings suggest that the developmental program of PLC SST neurons would define not only when these neurons *mature*, but also when they are *most susceptible* to biological and environmental modification. SST-expressing neurons in the PFC have been implicated in several models of affective behaviors in adult rodents, including fear conditioning (21), social fear (20), negative affect following stress (45), and substance use (23). Limited work has probed these neurons during similar behaviors in adolescent rodents, though neuropsychiatric illnesses often emerge during this period (1). We have previously shown that binge alcohol consumption disrupts firing properties of PLC SST neurons in an age-dependent manner, possibly reflecting a consequence of this extended period of plasticity. One potential mechanism for this is the expression of different ion channels in PLC SST neurons across ages, a mechanism which is supported by our changes in firing properties seen here. Our findings suggest that in addition to adolescence being a period during which the overall cortical circuit is being refined, critical components of the inhibitory microcircuit are themselves still under construction. Future work should determine which molecular signals regulate this prolonged developmental program and whether manipulating its timing alters adolescent vulnerability or resilience.

## Conclusions

Together, our findings reveal that SST neurons in PFC (higher-order association cortex) follow a fundamentally distinct developmental program from those in S1BF (somatosensory cortex). Across structural, intrinsic, and synaptic domains, prefrontal SST neurons continue to remodel throughout adolescence while homologous sensory circuits have largely stabilized. These results identify a genetically defined inhibitory microcircuit as a cellular substrate for prolonged prefrontal plasticity, providing a mechanistic framework for both the adaptability of adolescent cognition and the heightened vulnerability of the PFC to environmental and pathological insults during this developmental period. By defining when, and in which neurons, PFC inhibitory circuits remain plastic, we provide a foundation for understanding why adolescence represents a uniquely sensitive period for disorders involving executive function, affect regulation, and substance use.

## Acknowledgements

This study was supported by the National Institutes of Health [P50AA017823 (N.C.), R01AA029403 (N.C), R01AA031472 (N.C. and N.Z); T32GM154124 and F31AA030455 (A.R.S)]. The content is solely the responsibility of the authors and does not necessarily represent the official views of the National Institutes of Health.

## Materials and Methods

### Animals

Male and female homozygous SST-IRES-Cre (stock #013044), Ai9 (stock #007909), Ai32 (stock #024109), and mononucleotide repeated frameshift (MORF3; stock #035403) mice were purchased from The Jackson Laboratory for in-house breeding of hemizygous SST-Cre:Ai9, SST-Cre:Ai32, and SST-Morf3 offspring. Mice were maintained in a temperature- and humidity-controlled room (lights on at 7:00am). Age of pups were determined according to The Jackson Laboratory’s Mice Appearance by Age with PND 0 indicating the date of birth. Electrophysiology experiments were performed at 4 different age groups, with allowed age range in parentheses, to capture multiple stages of postnatal development: PND 14 (PND 13-16; pups were checked for eye opening prior to experimentation), PND 28 (PND 26-30), PND 42 (PND 40-44), and PND 56 (PND 54-58).

#### Immunohistochemistry

SST-Cre:MORF3 mice were bred to sparsely but fully label morphology of Cre-expressing cells (46) at PND 28 and PND 56. Details of immunohistochemistry are available in the **Supplementary Methods.**

#### Imaging and Morphology Analysis

*Imaging:* Images were taken as z-stacks with Leica STELLARIS 5 confocal microscope and were imaged at 20X magnification. Z-stacks were collapsed into a 2-dimensional image for analysis. 1-2 neurons were imaged per region per mouse.

*Sholl analysis:* dendritic intersections as a function of radius from the soma in ImageJ (National Institute of Health, Bethesda, MD, United States) using the Neuroanatomy plugin, blinded to the age cohort that the slice was from (47). The SNT toolbox of ImageJ allows semi-automated tracing of neuronal morphology and was used here to outline the soma and neuronal extensions from the soma using minimal individual pathways (47). The soma was excluded from the analysis to prevent it being quantified as multiple intersections.

#### Electrophysiology

Mice were transferred from the vivarium to the laboratory and allowed to acclimate for one hour prior to slicing. Slicing occurred at ZT 03:00 (+/- 1 hour), consistent with published experiments. Whole-cell patch-clamp electrophysiology was performed as previously described (48) at each of four age points. Mice were anesthetized with inhaled isoflurane and rapidly decapitated. Brains were rapidly extracted (<1 minute) and placed immediately in ice-cold N-methyl-D-glucamine (NMDG) solution: (in mM) 93 NMDG, 2.5 KCl, 1.2 NaH2PO4, 30 NaHCO3, 20 HEPES, 25 dextrose, 5 ascorbic acid, 2 thiourea, 3 sodium pyruvate, 10 MgSO4-7H2O, 0.5 CaCl2-2H2O, 306-310mOsm, pH 7.4 (49). Coronal sections containing the PL or S1BF cortex were prepared with a Compresstome vibrating microtome (Precisionary Instruments). Slices rested in oxygenated NMDG cutting solution (31°C) for a maximum of 8 minutes then were transferred to oxygenated artificial cerebrospinal fluid (aCSF) at 31°C, consisting of (in mM): 124 NaCl, 4.0 KCl, 1.2 MgSO4-7H2O, 2.0CaCl2-2H2O, 1 NaH2PO4, 25.95 NaHCO3, 9.99 dextrose, 305-308mOsmo. Slices rested in oxygenated aCSF for at least 1 hour prior to transfer to a submerged chamber for experiments. In the submerged chamber, slices were perfused with heated, oxygenated aCSF at a rate of 2mL/min. For optogenetic circuit mapping experiments, 3mM kynurenic acid (Sigma-Aldrich K3375) was added to the recording aCSF.

Recording electrodes (3-5 MΩ) were pulled from thin-walled borosilicate glass capillaries using a Narishige PC-100 Vertical Puller. Electrodes for all experiments were filled with a potassium-gluconate based recording solution (in mM): 135 potassium gluconic acid, 5 NaCl, 2 MgCl2-6H2O, 10 HEPES, 0.6 EGTA, 4 Na2ATP, 0.4 Na2-GTP, 287-290 mOsm, pH 7.35. Neurons were discarded from analysis if they had a resting membrane potential >-45mV, tonically fired action potentials, or had an access resistance exceeding 40 MΩ at the end of the experiment. A maximum of 3 cells per region were recorded from each mouse. Signals were digitized at 10 kHz and filtered at 4 kHz with a MultiClamp 700B amplifier. Recordings were analyzed using Clampfit 10.7 software (Molecular Devices, Sunnyvale, CA, United States).

### Intrinsic Excitability and Membrane Properties

In SST-Ai9 mice, SST-expressing cells in superficial layers of PL or S1BF cortex were identified by the presence of tdTomato fluorescence using 565nm LED excitation. Neurons expressing tdTomato which had a rheobase over 100pA or membrane resistance below 150MΩ, i.e. quasi fast-spiking non-Martinotti type SST neurons (50), were identified as statistical outliers so these cells were removed from all analyses.

Current-clamp experiments to assess the intrinsic excitability and properties of SST cells in both brain regions throughout development were performed. Following 5 minutes of baseline recording to allow the internal to diffuse in the cell, a depolarizing ramp current (0-100pA over 1 second) was injected into the cell to determine the rheobase. Current-induced action potential firing was assessed using discrete steps of hyperpolarizing to depolarizing current (−100 to 200pA in 10pA steps, each step lasting 250ms). Experiments were performed at each neuron’s resting membrane potential (RMP) then repeated at a common holding potential of –70mV. A description of the membrane properties assessed is available in the **Supplementary Methods**.

### Optogenetic Circuit Mapping

In SST-Ai32 mice, pyramidal neurons in superficial layers of PLC and S1BF were identified based on morphology and membrane properties (i.e. resistance <100MΩ or capacitance >75pF) as previously reported (21, 23) and patched with a potassium gluconate-based internal solution. Further details on optogenetic circuit mapping are available in the **Supplementary Methods**.

#### Data Analysis, Statistics, and Figure Preparation

Data were analyzed with 2-way ANOVAs (factors: age, sex), t-tests, and generalized linear mixed-effects models (GLME) where indicated. Except for GLME, statistics were run using GraphPad Prism 11.0. PCA was performed on 11 electrophysiological properties from 132 neurons. Data were standardized prior to PCA. Principal components were selected by Parallel Analysis. Scores for PC1-3 were extracted and analyzed with a three-way ANOVA (factors: sex, age, cell region). GLME were run in MATLAB 2021b as: Outcome Variable ∼ 1 + Cell Region x Age x Sex + (1 | Mouse), where mouse represents a random effect to account for collecting multiple cells from one animal. Figures were prepared using GraphPad Prism 11.0 (San Diego, CA). Data are presented as the mean ± the standard error of the mean.

## Supplementary Methods

### Immunohistochemistry

At PND 28 or PND 56, mice were anesthetized with inhaled isoflurane and perfused transcardially with cold phosphate buffered saline (PBS) for 3 minutes followed by 4% paraformaldehyde (PFA) in PBS for 3 minutes. Brains were extracted and stored in PFA at 4°C overnight before being moved to PBS at 4°C until slicing, at maximum 1 week after perfusion. Slices containing the PFC and S1BF were generated at 40um thickness using a Leica vibratome. Slices were stored in PBS until staining, at maximum 2 weeks after slicing. Immunohistochemistry for V5 fluorescent tag (Fortis Life Sciences A190-120A) was performed to label tagged SST neurons (46). Briefly, slices were rinsed in PBS 3 times for 10 minutes each and then blocked in 5% normal goat serum in PBS+0.1% Triton X-100 (Sigma X100). Slices were rinsed again 3 times in PBS for 10 minutes per wash then incubated overnight in primary rabbit polyclonal anti-V5 antibody (1:1000 in blocking solution). The next day, slices were rinsed 3 times in PBS for 10 minutes per wash then incubated in secondary Alexa Fluor 647 goat anti rabbit secondary antibody (Invitrogen A32733; 1:500 in PBS+0.1% Triton X-100) for 4 hours at room temperature. After 3 rinses in PBS for 10 minutes per wash, slices incubated in DAPI (1:10,000) for 10 minutes. Slices were rinsed in PBS 3 times for 5 minutes per wash before mounting and coverslipping with ImmunoMount.

### Electrophysiology

*Membrane properties were assessed as follows*:

− Resting membrane potential – measured after 5 minutes of gapfree
− Action potential threshold – for the first action potential elicited in the rheobase ramp current protocol, the threshold was calculated as the point of inflection for dV/dt
− Rheobase – measured as the current injected at the action potential threshold
− Membrane resistance – calculated as the slope of the I-V relationship for hyperpolarizing current steps from –50 to 0pA
− Membrane time constant – calculated by fitting an exponential curve to the end of the voltage response to a –50pA current injection
− Membrane capacitance – calculated by dividing the time constant by the input resistance
− Fast afterhyperpolarization (fAHP) - in rheobase, the difference between the action potential threshold and the peak minimum voltage
− Afterdepolarization (ADP) - in rheobase, the difference between fAHP and the immediate depolarizing overshoot
− Voltage sag ratio – following a hyperpolarizing step of –100pA –100pA, calculated as the peak hyperpolarization minus the steady state membrane potential, normalized to the steady state membrane potential (51)

### Optogenetic circuit mapping

Pyramidal neurons were held at 0mV (15) while two 1ms flashes of 470nm blue light (Cool LED, Traverse City, MI, United States) were delivered to evoke SST-mediated inhibitory postsynaptic currents (IPSCs). The flashes were delivered across interstimulus intervals from 100-1000 ms. Paired pulse ratio was calculated by dividing the peak of the second evoked IPSC by the peak of the first evoked IPSC, both normalized to the holding current. Properties of IPSC decay were measured from the first IPSC evoked at the 1000ms interstimulus interval. Time constant was measured from the peak of this first current to its return to baseline. For area under the curve, the baseline was adjusted to 0pA and area was measured across a standardized amount of time surrounding the first IPSC.

SST neurons in SST-Ai32 mice were also patched to confirm that optogenetic stimulation reliably evokes action potentials from SST neurons. SST neurons were held in current clamp and 470nm stimulation was delivered across various frequencies (5-40Hz).

## Supplementary Figures

**Supplementary Figure 1.**
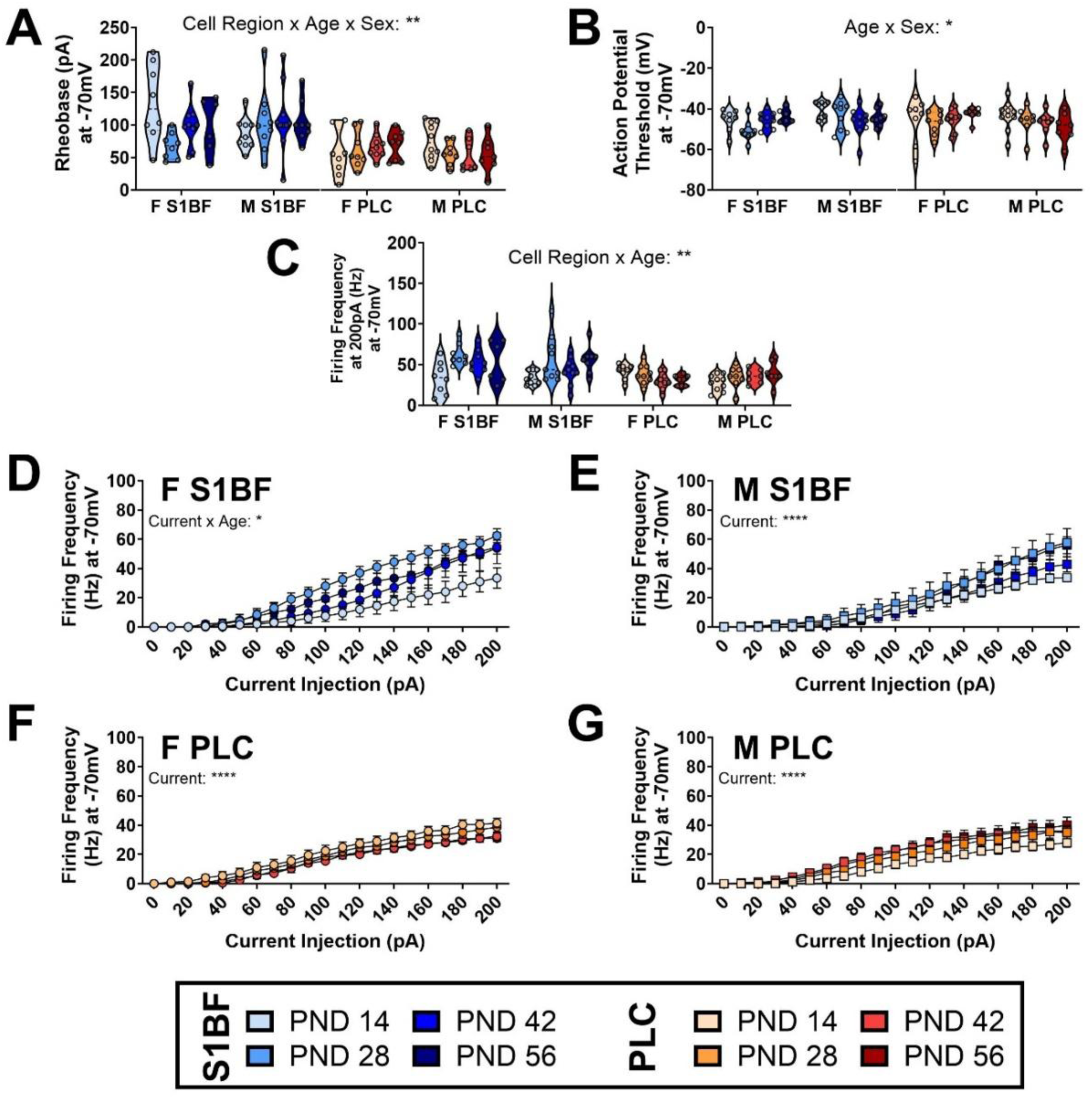
Excitability Properties of SST neurons at a common holding potential of –70mV. (A) The interaction between cell region, age, and sex was a significant predictor of rheobase of SST neurons while held at –70mV. (B) The interaction between age and sex was a significant predictor of action potential threshold of SST neurons held at –70mV. (C) The interaction between cell region and age significantly predicted peak firing frequency when SST neurons were held at -70mV. (D-G) There was a significant interaction between age and current on current-induced firing in F S1BF SST neurons but no other populations.

**Supplementary Figure 2.**
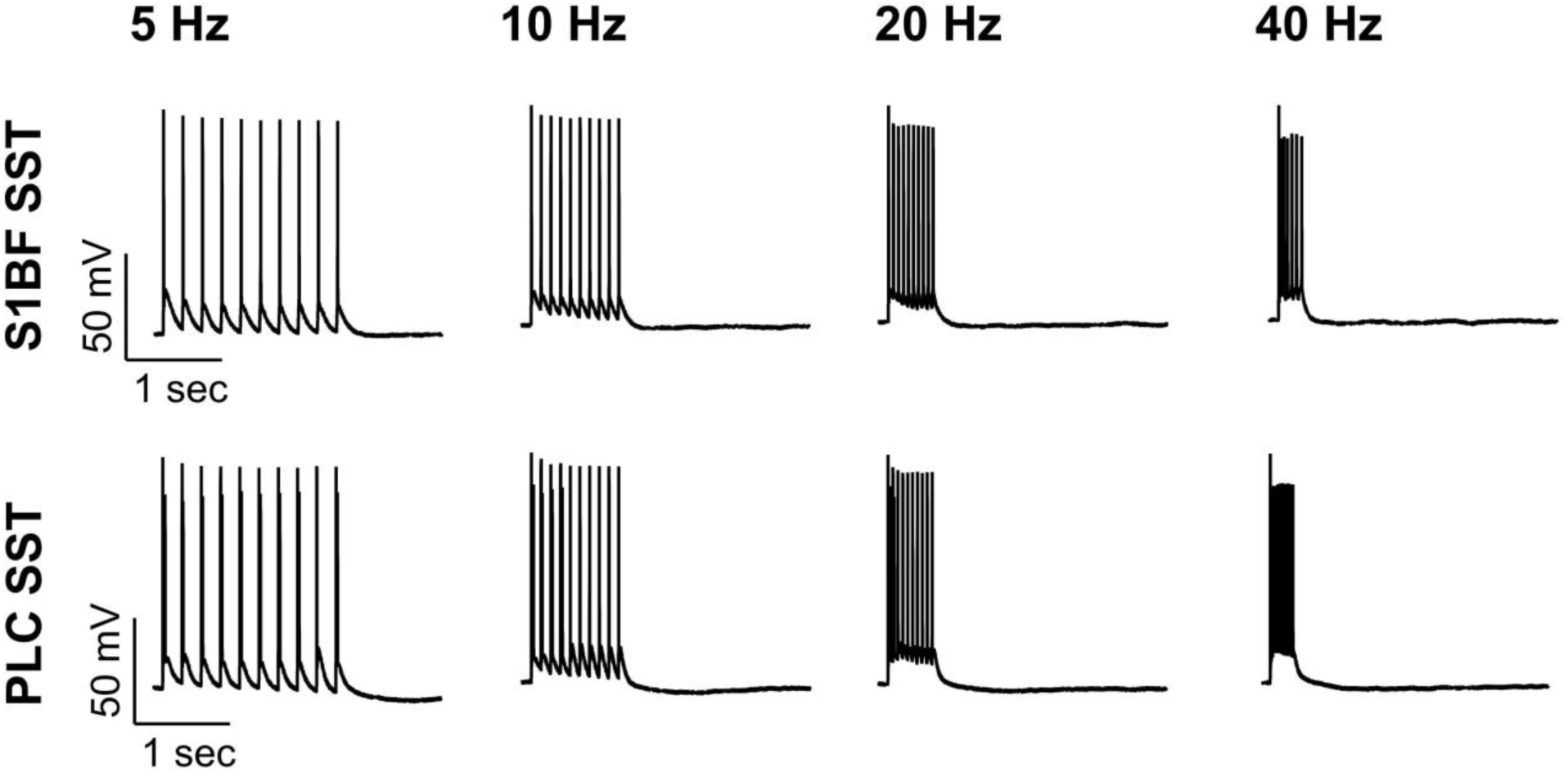
Representative traces showing that SST neurons in S1BF and PLC of SST-Ai32 mice robustly fire in response to 470nm light. Traces are taken from a PND14 mouse.

**Supplementary Figure 3.**
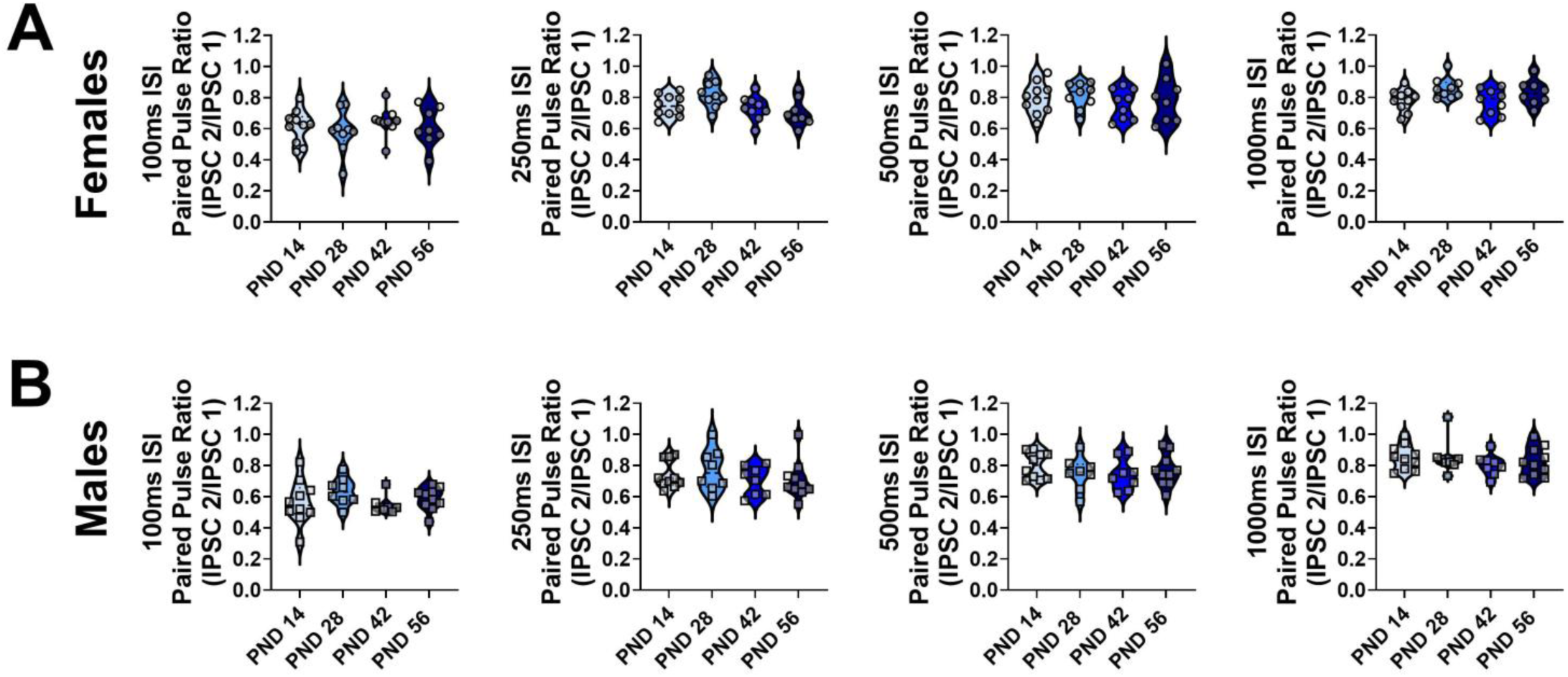
Paired pulse ratio of evoked IPSCs onto pyramidal neurons in S1BF of SST-Ai32 mice. (A) Paired pulse ratio across interstimulus intervals in female mice. (B) Paired pulse ratio across interstimulus intervals in male mice.

**Supplementary Figure 4.**
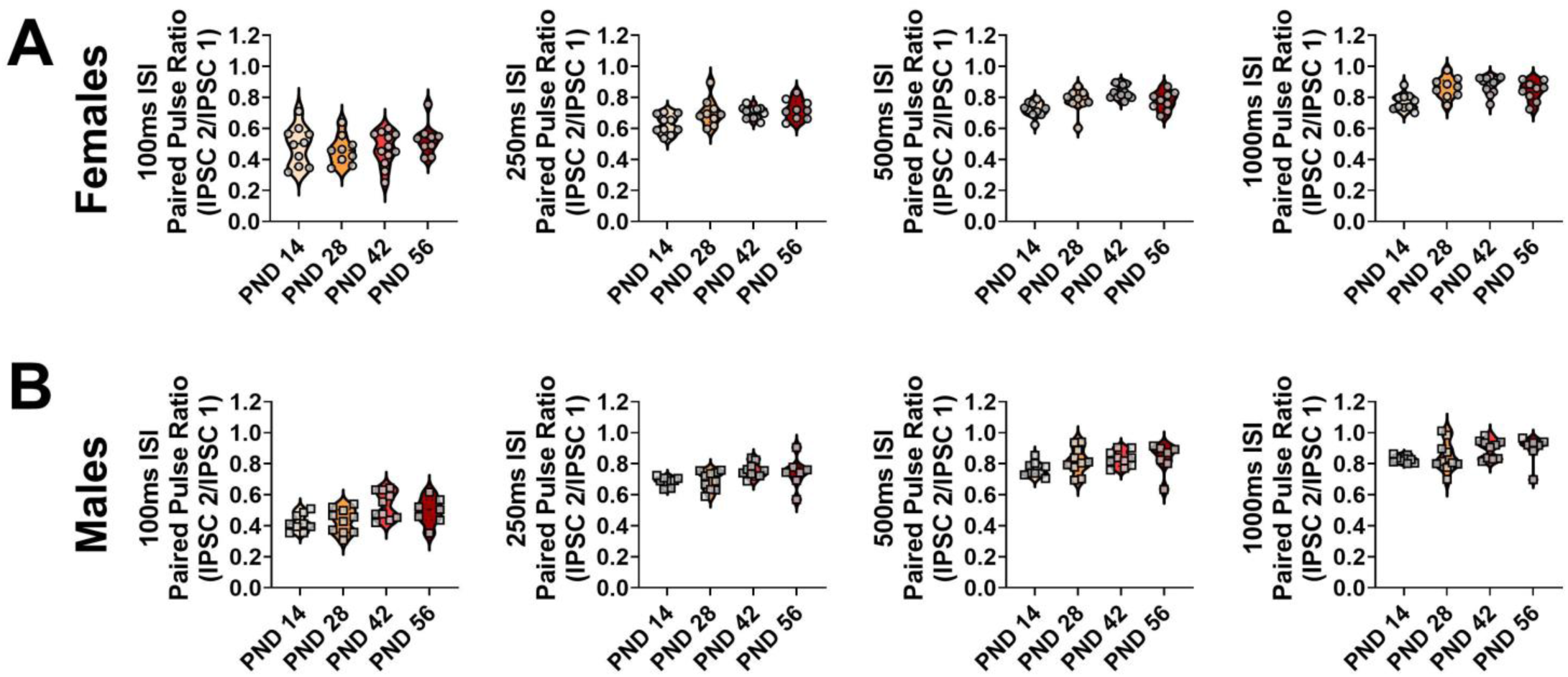
Paired pulse ratio of evoked IPSCs onto pyramidal neurons in PLC of SST-Ai32 mice. (A) Paired pulse ratio across interstimulus intervals in female mice. (B) Paired pulse ratio across interstimulus intervals in male mice.

